# Do Candidate Genes Affect the Brain’s White Matter Microstructure? Large-Scale Evaluation of 6,165 Diffusion MRI Scans

**DOI:** 10.1101/107987

**Authors:** Neda Jahanshad, Habib Ganjgahi, Janita Bralten, Anouk den Braber, Joshua Faskowitz, Annchen R Knodt, Hervé Lemaitre, Talia M Nir, Binish Patel, Stuart Richie, Emma Sprooten, Martine Hoogman, Kimm van Hulzen, Artemis Zavaliangos-Petropulu, Marcel P Zwiers, Laura Almasy, Mark E Bastin, Matt A Bernstein, John Blangero, Joanne Curran, Ian J Deary, Greig I de Zubicary, Ravi Duggirala, Simon E Fisher, Barbara Franke, Peter Fox, David Goldman, Asta K Haberg, Ahmad Hariri, L Elliot Hong, Matt Huentelman, Nicholas G Martin, Jean-Luc Martinot, Andrew McIntosh, Katie L McMahon, Sarah E Medland, Braxton D Mitchell, Susana Muñoz Maniega, Rene L Olvera, Jaap Oosterlaan, Charles Peterson, Natalie Royle, Andrew J Saykin, Gunter Schumann, John Starr, Elliot A Stein, Jessika Sussmann, Maria del C. Valdés Hernández, Dennis van’t Ent, Joanna M Wardlaw, Michael W Weiner, Douglas E Williamson, Anderson M Winkler, Margaret J Wright, Yihong Yang, Paul M Thompson, David C Glahn, Thomas E Nichols, Peter Kochunov

## Abstract

Susceptibility genes for psychiatric and neurological disorders - including *APOE, BDNF, CLU,CNTNAP2, COMT, DISC1, DTNBP1, ErbB4, HFE, NRG1, NTKR3*, and *ZNF804A* - have been reported to affect white matter (WM) microstructure in the healthy human brain, as assessed through diffusion tensor imaging (DTI). However, effects of single nucleotide polymorphisms (SNPs) in these genes explain only a small fraction of the overall variance and are challenging to detect reliably in single cohort studies. To date, few studies have evaluated the reproducibility of these results. As part of the ENIGMA-DTI consortium, we pooled regional fractional anisotropy (FA) measures for 6,165 subjects (CEU ancestry N=4,458) from 11 cohorts worldwide to evaluate effects of 15 candidate SNPs by examining their associations with WM microstructure. Additive association tests were conducted for each SNP. We used several meta-analytic and mega-analytic designs, and we evaluated regions of interest at multiple granularity levels. The ENIGMA-DTI protocol was able to detect single-cohort findings as originally reported. Even so, in this very large sample, no significant associations remained after multiple-testing correction for the 15 SNPs investigated. Suggestive associations (1.3×10^-4^ < p < 0.05, uncorrected) were found for *BDNF, COMT*, and *ZNF804A* in specific tracts. Meta-and mega-analyses revealed similar findings. Regardless of the approach, the previously reported candidate SNPs did not show significant associations with WM microstructure in this largest genetic study of DTI to date; the negative findings are likely not due to insufficient power. Genome-wide studies, involving large-scale meta-analyses, may help to discover SNPs robustly influencing WM microstructure.

## Introduction

Population and family studies revealed a key role for genetics in neuropsychiatric disorders. Over the last decade, genome-wide association studies (GWAS) that search for genetic risk factors underlying disease have successfully pinpointed common genetic variants found more frequently in affected than unaffected individuals. These genome-wide searches, which consider evidence for association across the genome, use a statistical correction to account for the number of tests performed. In theory, focused tests of candidate risk factors may provide a more efficient evaluation of mechanistic processes that affect gross anatomy and white matter microstructure.

We do not generally know the mechanisms by which genetic risk variants impact brain structure and function. The field of neuroimaging genetics may expedite such discoveries. Genetic factors influence variations in brain structure and organization, as measured with magnetic resonance imaging (MRI), and ultimately, genetic variations may modulate brain function and risk for disease. Recently, large-scale consortia −− including the Enhancing Neuro Imaging Genetics through Meta Analysis (ENIGMA) Consortium and Cohorts for Heart and Aging Research in Genomic Epidemiology Consortium (CHARGE) −− have identified consistent neuroanatomical associations with specific genetic variants. These collaborative approaches boost statistical power by assessing tens of thousands of people, and yield more accurate estimation of effect sizes ^1–4^.

Before the formation of these large-scale consortia, imaging genetics focused on mapping the effects of candidate SNPs and genes from the neuropsychiatric literature in the brains of smaller more targeted samples - often with several hundred individuals. Independent replication of genetic effects on brain anatomy is challenging, as common genetic variations exert small effects, accounting for <1% of the variance. The sample sizes of typical MRI studies do not have sufficient power to reliably detect these effects ^5,6^ and are more susceptible to false positive findings. However, without replication, true effects cannot be confirmed, and time and money may be wasted following up false positives. Several candidate SNPs influencing brain volume have been reported, however, most findings have not been replicated in large-scale consortium work: the majority of the candidates did not show even marginal effects in one study ^4^. These replication failures indicate that the initial reported effects may have been inflated, due to the ‘winner’s curse’ phenomenon ^7^. However, several other reasons for replication failures have been proposed. It could be argued that the meta-analytic approach typically adopted by GWAS consortia was limiting the possible power, or that minor methodological considerations in the imaging protocols were having a dramatic effect. The recent discovery of multiple significantly associated SNPs that account for less than 1% of the variance in regional brain volumes makes this less likely ^4^. Here, we aim to investigate these possibilities in the most methodologically comprehensive imaging genetics study to date examining candidate genes on brain structure. We analyzed over 6,000 scans from 11 participating cohorts from the Americas, Europe, and Australia.

As shown by ENIGMA and other consortia, MRI can help to discover genetic variants that influence brain and substructure volumes. Genetic studies of diffusion MRI can examine how genetic variants impact white matter pathways. Diffusion tensor imaging (DTI) quantifies the microstructural properties of white matter and allows for network type reconstructions of the brain’s physical connections. DTI measures are altered in a wide range of diseases, including stroke ^8^, Alzheimer’s disease (AD) ^9,10^, schizophrenia ^11,12^, bipolar and mood disorders ^13,14^, and addiction ^15,16^, among others. The heritability of many DTI measures has been established; our DTI group within ENIGMA has found these heritability estimates to be reliable and remarkably similar across several cohorts ^17–19^. As approximately half of the variability in many of these DTI measures is due to additive genetic effects, it is expected that genetic association studies will provide insight into the mechanisms, by which these connections are formed and maintained.

Several published works have already proposed genetic associations between candidate SNPs and white matter microstructure as defined by DTI scans (See Table 1). Independent replication of these results, however, has been limited. The initial findings were generally made in samples of 100-1000, large for an imaging study, but arguably too small to robustly estimate effect sizes. Similar to several early candidate SNP effects having been disputed in the psychiatric literature ^20,21^, the effects of candidate variants on brain structure have become increasingly controversial ^22^. ENIGMA’s subcortical volume GWAS identified robust associations accounting for as little as 0.5% of the variance in volumes of specific structures, but the majority of SNPs identified as influencing psychiatric disorders do not reach nominal levels of significance ^4^.

**Table 1:**
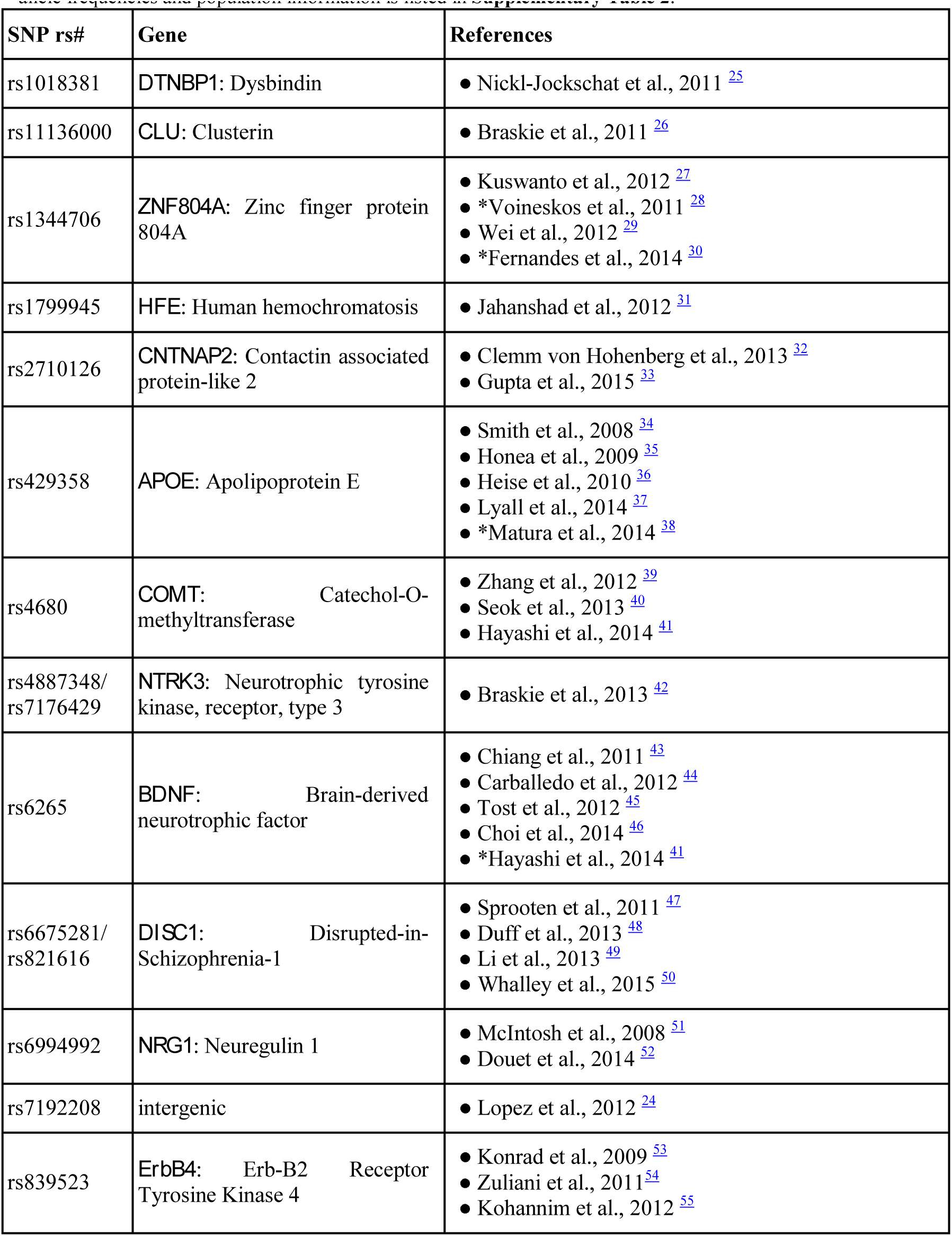
List of candidate SNPs with their corresponding genes. These SNPs have been investigated to find associations with FA derived from diffusion tensor imaging (DTI). References to associations or lack of associations are listed in the last column. Published null findings are marked with a ‘*’. Additional information including minor allele frequencies and population information is listed in **Supplementary Table 2**.

Here, we investigated the effects of candidate variants on the microstructure of white matter pathways as defined by the most common diffusion MRI measure, fractional anisotropy (FA). In these analyses, FA was averaged across the full WM skeleton as well as specific tracts using harmonized protocols described in our prior work ^17–19^. We assembled a sample of 6,165 individuals (4,458 of them of Caucasian (CEU) ancestry) with DTI and genetic information provided by eleven cohorts. We limited the SNPs considered to those previously reported to show associations with FA. While most of the SNPs we targeted were originally reported as risk factors in the neuropsychiatric literature, one SNP was first identified in a genome-wide analysis of DTI-FA traits as being in a promising genetic locus (though it was not shown to be genome-wide significant). We included SNPs in the candidate genes *APOE, BDNF, CLU, CNTNAP2, COMT,DISC1, DTNBP1, ErbB4, HFE, NRG1, NTKR3*, and *ZNF804A*, summarized in Table 1.

We assessed the association of these variants in the standard meta-analytical framework used for GWAS. We also determined whether the power to detect the genetic effects would be improved in a mega-analytical framework, where the statistical tests are computed on a pooled data set. In addition, we examined the effect within the samples in which the associations had originally been reported. This allowed us to determine whether the harmonized image processing approach taken within ENIGMA would maintain the nominal association within that sample.

Independent testing of multiple brain regions of interest may impose an unnecessarily strict multiple comparisons correction requirement on the results, if the average FA across the full brain has similar effect sizes for the individual SNPs as the regional measures. Therefore, as a follow-up analysis, we made statistical inferences across various groupings of the regions of interest to determine if effects would be stronger had we, for example, analyzed bilateral regions separately, or only analyzed the corpus callosum as a whole, rather than including the genu, body, and splenium, separately.

We hypothesized that the reported findings may be limited to the discovery samples, and may not generalize to other cohorts, reinforcing the need for replication in imaging genetics. We further hypothesized that the impact of methodological differences in image processing on the association would be limited, so that associations reported in the original findings would be present in the phenotypes produced using the harmonized ENIGMA protocols, and the effect sizes would be similar for each SNP between meta-and mega-analyses of cohort effects. Our study corroborates the inconsistencies in the current literature and provides ample evidence for the need to continue unbiased genome-wide searches with quantitative brain measures.

## Materials and Methods

Our study aimed to determine whether candidate SNPs, that were previously reported to have an association with white matter microstructure (as measured by DTI fractional anisotropy, FA), can be replicated in a large DTI-based imaging genetics study. In the following sections, we describe: (2.1) the selection process for SNPs we focused on in this study; (2.2) demographic and imaging parameters of the cohorts included in the analysis; (2.3) a brief overview of the previously described harmonized image processing performed as part of ENIGMA-DTI; (2.4) the genetic associations and the meta-and mega-statistical approaches; (2.5) multiple comparisons correction considerations taken in this work.

### 2.1 Candidate SNP selection

We identified the SNPs considered here through a literature search, when this project was initiated in June 2014, setting a filter to find those SNPs associated with FA that had been selected because of (1) their ‘known’ links with brain-related disorders or (2) those that had been discovered through genome-wide screens of diffusion-weighted imaging measures. This search identified fourteen SNPs within twelve genes (*APOE, BDNF, CLU, CNTNAP2, COMT, DISC1,DTNBP1, ErbB4, HFE, NRG1, NTKR3, ZNF804A*) that had previously been identified as candidate genes within the psychiatric literature (See Table 2). GWASs for DTI-FA have also been performed ^23,24^, but no results have met statistical thresholds for genome-wide significance. To determine if the strongest hit from these analyses would reach genome-wide significance if the overall sample size was dramatically increased, this variant was also included as a putative candidate. All of these SNPs are common (present in >5% of the population) and have been previously associated with normal variance in FA values in healthy individuals, in at least one published study.

**Table 2:**
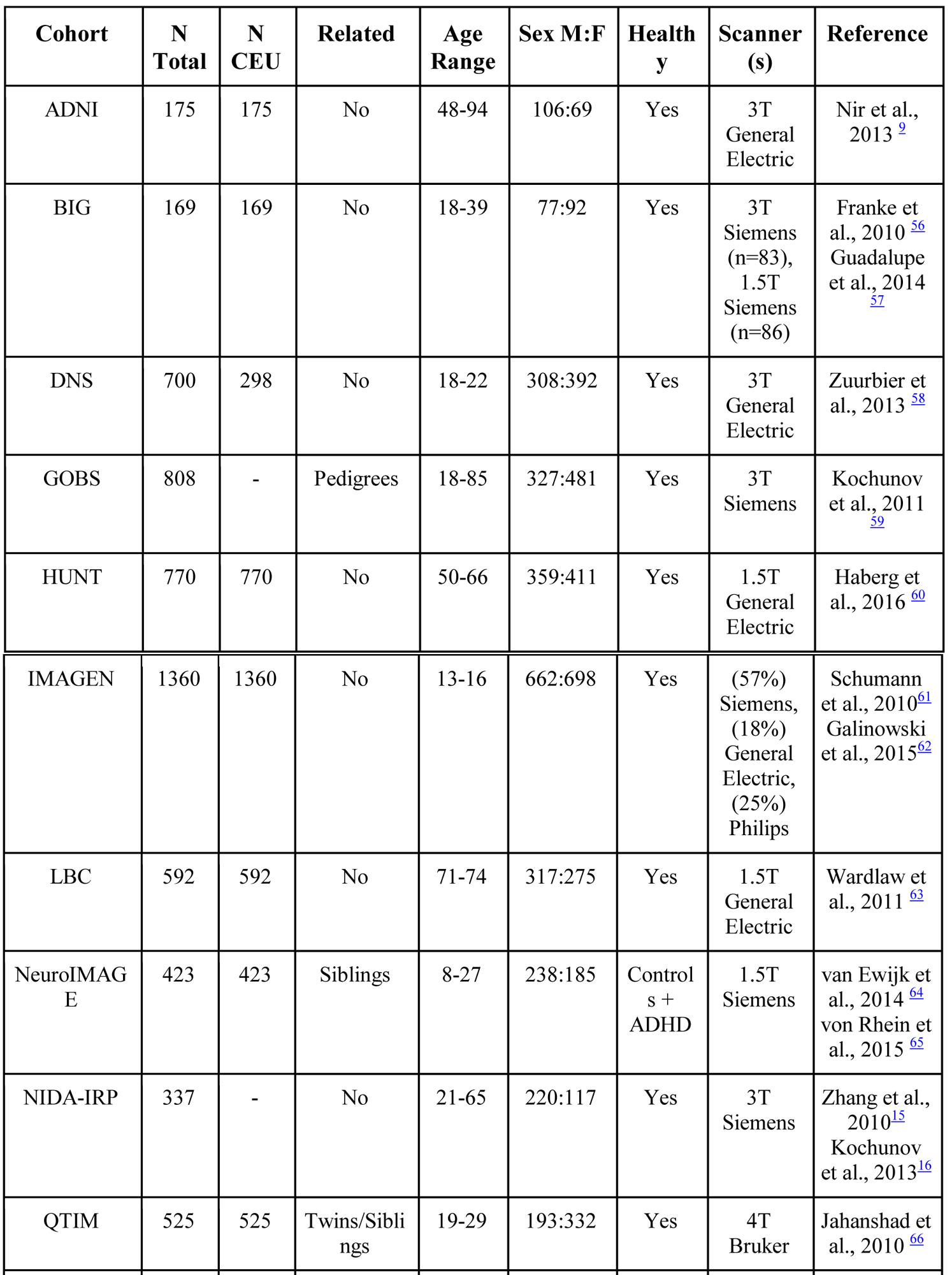
The demographic breakdown of the 11 cohorts included in the study is listed below: ADNI: Alzheimer’s Disease Neuroimaging Initiative (healthy only subsample used here); BIG: Brain Imaging Genetics; DNS: Duke Neurogenetics Study; GOBS: Genetics of Brain Structure and Function; HUNT: The Nord-Trøndelag Health Study; IMAGEN: European research project investigating mental health and risk taking behaviour in teenagers; LBC: Lothian Birth Cohort; NeuroIMAGE: integrated DNA-cognition-MRI-phenotype project with the aim to identify cognitive, neural and genetic underpinnings of ADHD; NIDA-IRP: National Institute on Drug Abuse, Intramural Research Program; QTIM: Queensland Twin IMaging Study; TAOS: Teen Alcohol Outcomes Study.

### 2.2 Cohorts included

Eleven cohorts were included in this study, leading to a dataset totaling 6,165 individuals (ages: 11-85). The imaging genetics analysis tool, Sequential Oligogenic Linkage Analysis Routines (SOLAR)-Eclipse software package (http://www.nitrc.org/projects/se_linux), was used to create kinship matrices for each cohort. Cohorts consisted of various twin, family, and unrelated individuals, mainly of European descent. Cohorts of individuals from mixed ethnic populations in the United States, including Latinos and African Americans as well as Caucasians, were also included. Genetic imputation was performed according to the ENIGMA Imputation cookbook (http://enigma.ini.usc.edu/wp-content/uploads/2012/07/ENIGMA2_1KGP_cookbook_v3.pdf) for most sites. The genetic variability of the individuals in the samples was compared to that of known population cohorts through multidimensional scaling (MDS); the first two components for each individual were plotted against known ethnic samples from the HapMap3 efforts. To avoid effects of population stratification, analyses were performed in the largest ethnic group (corresponding to 4,458 from the CEU population) and subsequently in all individuals. All meta-and mega-analytical approaches were used to estimate overall genetic associations in both cases.

### 2.3 ENIGMA-DTI processing

We used the ENIGMA-DTI protocols for multi-site processing and extraction of tract-wise average FA values, as described in our prior work ^17–19^. Briefly, FA images from all subjects were non-linearly registered to the ENIGMA-DTI target FA map. This target was created from the FA images of four participating studies and has been shown to provide stable results for children and adult cohorts, as previously described ^17^. Registration algorithms were allowed to vary to ensure optimal alignment per cohort, yet FSL’s *fnirt*^69^ was suggested as a default registration, with an optional additional script provided using ANTS (http://stnava.github.io/ANTs/) registration tools. The data were then processed using the tract-based spatial statistics (TBSS) analytic method ^70^ modified to project individual FA values on the ENIGMA-DTI skeleton. Following the extraction of the skeletonized white matter and projection of individual FA values, ENIGMA tract-wise regions of interest, derived from the Johns Hopkins University (JHU) white matter parcellation atlas ^71^, were transferred to extract the mean FA across the full skeleton and average FA values for a total of 25 (partially overlapping) regions [**SI Figure 1 and SI Table 1**] found to be reliably heritable in our prior work ^17–19^. The protocol, quality control scripts, target template, skeleton mask, source code, and executables are publicly available at https://www.nitrc.org/projects/enigma_dti/ or http://enigma.ini.usc.edu/ongoing/dti-working-group/.

### 2.4 Site-level and pooled genetic associations

We used several statistical models in this analysis; full details can be found in the **Supplementary Text**. Here, we briefly describe the models and the covariates used.

Regressions were run in the full white matter average FA and for each of 21 unique bilateral regional measures, as outlined by the ENIGMA-DTI protocols in Section 2.3. Abbreviations and full tract names are listed in **SI Table 1**. The protocol also contained 3 additional measures of FA combined for larger tracts, including the entire corpus callosum (3 original regions: genu, body splenium), internal capsule (3 original regions: anterior limb, posterior limb, and retrolenticular), and *corona radiata* (3 original regions: anterior, posterior, superior); this totals 25 (4 non-unique) measures of FA being tested. FA values were used as-is (raw and un-normalized), as well as after applying an inverse Gaussian normalization to the residuals after adjusting for effects of the covariates. Inverse-variance weighted meta-analysis was performed to pool resulting inferences on both sets, and mega-analyses were performed on the normalized measures.

Each SNP (coded in a dose-dependent, additive fashion with respect to the minor allele count 0,1, or 2) was regressed on the average FA in each ROI independently at the site level, and subsequently at the group level for mega-analysis. Linear models were run for all cohorts of unrelated individuals, and mixed effects models using the kinship matrix to model relatedness were used for all cohorts with related individuals. We used SOLAR-Eclipse (http://www.nitrc.org/projects/se_linux) to create kinship matrices for each individual cohort of related individuals after specifying familial relationships. All regressions were performed in the CEU sample only as well as the full cohort.

Several fixed covariates were used in all analyses, including age, sex, age-by-sex, age-squared, and age-squared-by-sex. Nonlinear effects of age (i.e. age-squared, and age-squared-by-sex) have been shown to impact FA ^59^ and were included for all cohorts. The first four components of the MDS analyses were also used as covariates for all groups. Two sites, NeuroIMAGE and BIG, added a covariate to model the two scanners at which the images were collected. NeuroIMAGE also added a covariate to model potential fixed differences between participants with ADHD and controls.

In addition to the breakdown of ROIs tested as part of the ENIGMA-DTI protocol, to assess the regional specificity of the associations, we tested various groupings of the ROIs. Bilateral measures were broken down into their lateralized regions, and results of those associations statistically pooled using Stouffer’s meta-analysis ^72^ were compared to the original groupings, which were further pooled and compared to the effects computed on the overall average FA; we aimed to pool regional estimates and therefore reduce the correction needed for ROIs. Methods and detailed results of this analysis can be found in the **Supplementary Text**.

### 2.5 Multiple comparisons correction

Multiple statistical tests performed across regions of the image can increase the chance of reporting a false positive finding at a given significance threshold, unless steps are taken to control for multiple comparisons. A Bonferroni correction threshold accounts for the number of variants tested; here 15 variants were tested, leading to a single trait correction threshold of 0.05/15 = 0.0033. However, as no SNPs reached levels of significance of *p* < 0.0033, we also report those that reached a nominal significance, defined here as those having an uncorrected *p*-value of *p* < 0.05. While neglecting that multiplicity increases false positive risk, with such negative significance tests, it can only strengthen the evidence of null effects.

## Results

No statistically significant findings were observed when data was pooled across studies, regardless of pooling method, although most SNPs were found to have similar directions of effect across many regions of interest (Figure 1).

**Figure 1.**
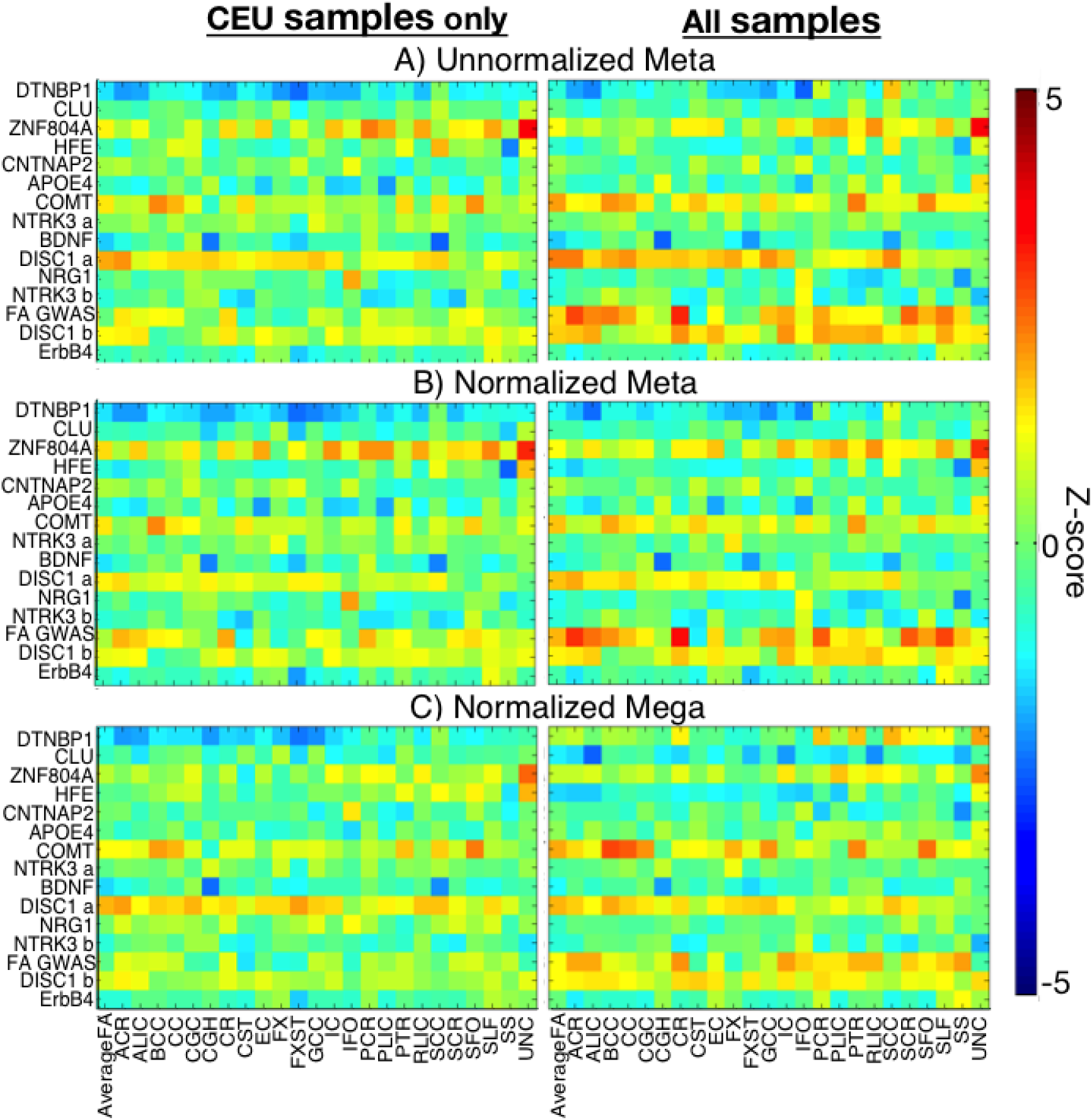
**Effects in people with European ancestry and all ancestries combined.** The normalized effect size is shown as the Z-score of the joint associations for each SNP on each regional measure (DTI measures are shown on the *x*-axis). The *left-most column* highlights the effect of the SNPs in the CEU subpopulation and the *right-most column* in all 6,165 individuals. The *top-most plots* (A) are the meta-analyzed results of the association tests that used a linear regression model (accounting for kinship) for each cohort on the variable of interest directly. The *middle row* (B) shows results of the same regression model on the residuals of the variable after removing the effects of covariates; the *bottom-most row* (C) is the result of the mega-analysis run on the residuals at every site. Some SNPs showed nominal significance (*p*<0.05) regardless of the model chosen, but no SNP survived multiple comparisons correction for the number of SNPs tested across ROIs. SNPs are labeled according to their gene name; two SNPs in *DISC1* and *NTRK3* were evaluated *DISC1*-a refers to rs6675281, *DISC1*-b to rs821616; *NTRK3*-a to rs4887348 and *NTRK3*-b to rs7176429.

The breakdown of SNP counts per cohort is highlighted in **SI Table 2**. The minor allele frequency (MAF) for the *dysbindin* SNP rs1018381 was low (<10%) for all CEU groups. GOBS and NIDA cohorts had a higher rs1018381 MAF, perhaps due to their more diverse ethnic composition. While all other SNPs had a MAF > 10% in the CEU population, the *DISC1* SNP rs6675281 had no homozygous carriers in GOBS, and the *HFE* SNP rs1799945 had no homozygotes in NIDA-IRP, both non-CEU cohorts.

As seen in Figure 2, correlations between the methods were strong in the CEU subpopulation; within the full sample, the mega-analyses revealed weaker correlations, likely due to the population heterogeneity, variations in the MDS component scale as well as SNP frequency, reiterating a known need for cautionary procedures, when combining multiple cohorts. Consistencies between the various meta-analytical approaches confirmed that the FA distributions with current sample sizes were sufficient to use raw-level statistics without inverse Gaussian normalization and parametric approaches.

**Figure 2.**
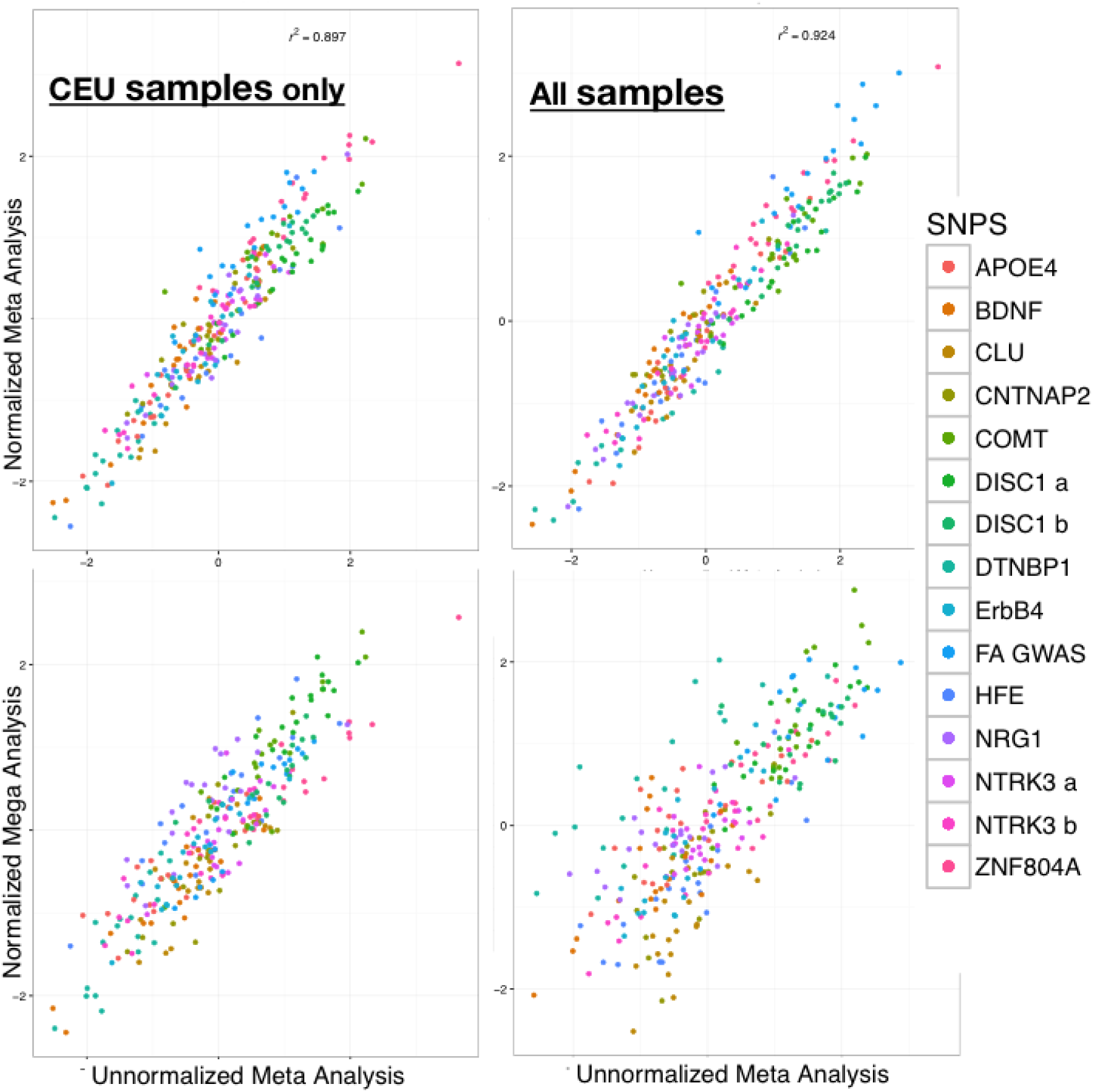
Correlations between the overall z-scores across the different methods are plotted. Normalized and non-normalized meta-analyses are very highly correlated for both the CEU subsample (r^2^=0.90) and the full sample (r^2^=0.92). The mega-analysis is also highly correlated for the CEU sample (r^2^=0.8). In the full sample, the mega-analyses revealed weaker correlations (r^2^=0.63). SNPs are labeled according to their gene name; two SNPs in *DISC1* and *NTRK3* were evaluated *DISC1*-a refers to rs6675281, *DISC1*-b to rs821616; *NTRK3*-a to rs4887348 and *NTRK3*-b to rs7176429.

Within the CEU sample, nominally significant (uncorrected *p*<0.05) associations were found with both meta-and the mega-pooling method for three polymorphisms: the first schizophrenia GWAS-significant SNP in *ZNF804A*, rs1344706, in the uncinate, *COMT* rs4680 in the body of the corpus callosum, and *BDNF* rs6265 in the splenium of the corpus callosum and the hippocampal portion of the cingulum. Forest plots for these findings are shown in **SI Figure 4**. The standard errors for the associations computed with mega-analysis were marginally larger than those computed with meta-analysis (shown by width of the markers in the Forest plots in **SI Figure 4**). In the plots, the size of the center square reflects the sample size of the cohort; these can be seen for raw-valued associations and those Gaussian-normalized before regression and joint statistics. Additional nominally significant associations were found with two of the three approaches (both meta, or mega and one meta); these included *HFE* rs1799945 in the sagittal stratum, *ZNF804A* rs1344706 in the SLF, PCR, PLIC, and RLIC, and *DISC1* rs6675281 in the anterior *corona radiata*. Nominal associations were also found for *DTNBP1* rs1018381, but as previously mentioned, MAFs were too low in the CEU samples for reliable analysis. Our analysis of pooling ROIs through Stouffer’s meta-analysis was similar to our individual ROI findings and did not reach statistical significance.

To establish whether the negative results were the result of insufficient sensitivity, we conducted a power analysis ^73,74^ using a t-test linear multiple regression design in G*Power 3.1 ^75^. Previous studies had found SNPs that explain 0.5-1% of the phenotypic variance; using these effect sizes with N=4,458 subjects, and 18 fixed effects (11 sites, five age− and sex-related covariates, and two site-specific dummy variables) with an alpha of 0.05/15/25=1.3×10^-4^ to reflect the 15 SNPs and search regions (if all 25 were independent), gives a power in the range of 81.5 to 99.8%. Thus, these negative results are unlikely to be explained by insufficient power.

## Discussion

We performed a targeted investigation of candidate SNPs implicated in disease risk that have been previously reported to also influence white matter microstructure. SNPs in multiple genes have been claimed to be associated with variation in the white matter microstructure of the brain as measured by FA from DTI. To test the validity of the associations, it is important to ensure that the findings are replicable across populations, and that sample sizes are large enough to detect statistically significant associations. While no single imaging site has the sample size needed, effective ways to harmonize analyses and pool existing data sets from across the world are needed to discover robust associations. Such multi-site, international studies allow discovery of genetic effects in populations around the world, and ensure that these associations are consistent in diverse groups of individuals. The diversity in this context is not genetic diversity in ancestry, which could lead to population stratification effects, but diversity in terms of various environmental and demographic aspects captured through global initiatives within the ENIGMA Consortium. Here, the statistical tests were performed centrally and on the full set of data at once, whereas in a harmonized meta-analytic framework statistics are conducted individually for each dataset and summary results are pooled across studies.

While some SNPs showed nominal associations with localized regions within the FA map, this study did not replicate the association of any of the 15 SNPs previously associated with FA in single cohort studies, suggesting that previously reported associations might not be robust or generalizable. Classic candidate genes long thought to play a significant role in the genetic basis of schizophrenia, including some of those evaluated here, such as *COMT* and *DISC1*, were recently re-evaluated in the largest genomic study of schizophrenia published to date ^20,21^. These traditionally studied genetic factors were found not to be as pertinent to the disease as initially hypothesized, and it was suggested that the original findings were perhaps driven by relatively small and underpowered studies. The situation for imaging genetics studies cannot be assumed to be any different from these studies of schizophrenia, or any other complex human traits, in that regard.

Our findings may potentially be discouraging to researchers whose studies focus on these candidate SNPs, but they do not rule out that effects of such SNPs might be observed in even larger sample sizes. Moreover, we targeted specific SNPs of interest within each candidate gene, and cannot discount potential contributions of other polymorphisms elsewhere in such genes, in regions of independent linkage disequilibrium. Indeed, the genetic architecture of these white matter traits may involve a broad range of common variants not yet identified, but identifiable with GWAS, as we have recently shown for structural MRI ^4^. This work shows that the individual associations reported in single cohorts are not reliably replicated in other cohorts. It also shows that meta-analysis approaches have similar power to mega-analysis. As none of the prior candidate SNPs showed widespread associations, we are unable to determine, if mega-analysis offers more power to detect *true* associations than meta-analysis. In heterogeneous cases, we found that meta-analytic approaches may provide more consistent results than mega-analysis, while both meta- and mega-analysis performed appropriately for cohorts from ethnically homogeneous populations.

This is the first large-scale multi-cohort investigation of white matter microstructure assessing multiple candidate SNPs in a statistically rigorous and extensive evaluation. Our results question the reliability of previous findings, despite the historical significance many of these genes may have presented. However, the impact of “winner’s curse” and publication bias need to be taken into account, when reading imaging genetics literature that does not include evidence for replication. This is not intended to negate previous findings, as there are also population considerations that could perhaps influence the degree and direction of genetic associations, such as age, sex, and environmental effects. In this analysis, all groups span a variety of age ranges from multiple countries around the world. Our aim was to identify generalized associations of previously highlighted candidate SNPs with white matter across *multiple* populations around the world, not to negate findings within any particular subpopulation, in which they were originally reported; subpopulation associations may be valid and are beyond the scope of this study. Instead, we advocate the importance of reproducing effects, and highlight the need for harmonizing analyses, showing that the ENIGMA-DTI processing method was able to maintain the SNP associations within the cohort in which they were first reported. We show that these historical candidate SNPs are not globally and reliably ones that influence brain microstructure, and instead suggest that to identify the key genetic factors, a large-scale GWAS is needed, as in the successful GWAS reported for MRI-based measures of brain volumes ^1–4^.

Neuroimaging genetics offers a non-invasive window to observe genetic influences on macro-scale brain structure. However, to fully understand the mechanisms behind any association, more costly and invasive paradigms are necessary. It is important to ensure the robustness and generalizability of the imaging association before investing in in-depth analysis of a gene or specific locus; otherwise, significant resources may be wasted. The ENIGMA-DTI working group is now pooling together a full-scale detailed genome-wide meta-analysis of all common genetic variants in the genome in over 10,000 individuals, to contribute to an unbiased search for common genetic variants that help to shape white matter microstructure in the general population.

## Acknowledgements

This work is funded by the NIH U54 Big Data to Knowledge (BD2K) Initiative under U54EB020403 (PI: Thompson) to support big data analytics, management and distribution of programs. Additional support was provided by R01EB015611 (PI: Kochunov). Individual cohort funding is provided by:

**ADNI**: Data used in preparation of this article were obtained from the Alzheimer’s Disease Neuroimaging Initiative (ADNI) database (adni.loni.usc.edu). As such, the investigators within the ADNI contributed to the design and implementation of ADNI and/or provided data but did not participate in analysis or writing of this report. A complete listing of ADNI investigators can be found at: http://adni.loni.usc.edu/wp-content/uploads/how_to_apply/ADNI_Acknowledgement_List.pdf Data collection and sharing for this project was funded by the Alzheimer’s Disease Neuroimaging Initiative (ADNI) (National Institutes of Health Grant U01 AG024904) and DOD ADNI (Department of Defense award number W81XWH-12-2-0012). ADNI is funded by the National Institute on Aging, the National Institute of Biomedical Imaging and Bioengineering, and through generous contributions from the following: Alzheimer’s Association; Alzheimer’s Drug Discovery Foundation; Araclon Biotech; BioClinica, Inc.; Biogen Idec Inc.; Bristol-Myers Squibb Company; Eisai Inc.; Elan Pharmaceuticals, Inc.; Eli Lilly and Company; EuroImmun; F. Hoffmann-La Roche Ltd and its affiliated company Genentech, Inc.; Fujirebio; GE Healthcare; IXICO Ltd.; Janssen Alzheimer Immunotherapy Research & Development, LLC.; Johnson & Johnson Pharmaceutical Research & Development LLC.; Medpace, Inc.; Merck & Co., Inc.; Meso Scale Diagnostics, LLC.; NeuroRx Research; Neurotrack Technologies; Novartis Pharmaceuticals Corporation; Pfizer Inc.; Piramal Imaging; Servier; Synarc Inc.; and Takeda Pharmaceutical Company. The Canadian Institutes of Health Research is providing funds to support ADNI clinical sites in Canada. Private sector contributions are facilitated by the Foundation for the National Institutes of Health (http://www.fnih.org). The grantee organization is the Northern California Institute for Research and Education, and the study is coordinated by the Alzheimer’s Disease Cooperative Study at the University of California, San Diego. ADNI data are disseminated by the Laboratory for Neuro Imaging at the University of Southern California.

**BIG**: This work makes use of the Brain Imaging Genetics (BIG) database, first established in Nijmegen, The Netherlands, in 2007. This resource is now part of Cognomics (http://www.cognomics.nl), a joint initiative by researchers of the Donders Centre for Cognitive Neuroimaging, the Human Genetics and Cognitive Neuroscience departments of the Radboud University Medical Centre and the Max Planck Institute for Psycholinguistics in Nijmegen. The Cognomics Initiative is supported by the participating departments and centres and by external grants: the Biobanking and Biomolecular Resources Research Infrastructure (Netherlands) (BBMRI-NL), the Hersenstichting Nederland, and the Netherlands Organisation for Scientific Research. BIG also received funding from the European Community’s Seventh Framework Programme (FP7/2007 – 2013) under grant agreement number 602450 (IMAGEMEND).

**DNS**: Duke University and NIDA grant DA033369, R01-AG049789 and R01-DA031579

**GOBS**: The GOBS study (PI DG and JB) was supported by the National Institute of Mental Health Grants MH0708143 (Principal Investigator [PI]: DCG), MH078111 (PI: JB), and MH083824 (PI: DCG & JB).

**HUNT**: Liaison Committee between the Central Norway Regional Health Authority (RHA) and the Norwegian University of Science and Technology. The authors would like to thank The Nord-Trøndelag Health Study (The HUNT Study) for all their help. The HUNT Study is a collaboration between HUNT Research Centre (Faculty of Medicine, Norwegian University of Science and Technology NTNU), Nord-Trøndelag County Council, Central Norway Health Authority, and the Norwegian Institute of Public Health.

**IMAGEN**: IMAGEN was supported by the European Union-funded FP6 Integrated Project IMAGEN (Reinforcement- related behaviour in normal brain function and psychopathology) (LSHM-CT-2007-037286), the FP7 projects IMAGEMEND (602450) and MATRICS (603016), and the Innovative Medicine Initiative Project EU-AIMS (115300-2), the Medical Research Council Programme Grant “Developmental pathways into adolescent substance abuse” (93558), as well as the NIHR-biomedical Research Center "Mental Health". Further support was provided by the Swedish Research Council FORMAS and the German Federal Ministry for Education and Research BMBF (eMED SysAlc 01ZX1311A; Forschungsnetz AERIAL; 1EV0711).

**LBC1936**: We thank the members of the LBC1936 who participated in this study, radiographers at the Brain Research Imaging Centre (BRIC; http://www.bric.ed.ac.uk), and the LBC1936 research associates who collected and entered some of the data used in this manuscript. This research was supported by Research into Ageing and continues as part of The Disconnected Mind project, funded by Age UK. MRI acquisition and analyses were conducted at BRIC, Neuroimaging Sciences, University of Edinburgh which is part of SINAPSE (Scottish Imaging Network—A Platform for Scientific Excellence) collaboration (http://www.sinapse.ac.uk) funded by the Scottish Funding Council and the Chief Scientist Office. This work was undertaken within The University of Edinburgh Centre for Cognitive Ageing and Cognitive Epidemiology (http://www.ccace.ed.ac.uk), part of the cross council Lifelong Health and Wellbeing Initiative (MR/K026992/1), for which funding from the BBSRC and MRC is gratefully acknowledged.

**NeuroIMAGE**: This work was supported by National Institutes of Health (NIH) grant R01MH62873 (Stephen V. Faraone), Netherlands Organisation for Scientific Research (NWO) Large Investment Grant 1750102007010 (Jan K.Buitelaar) and grants from Radboud University Nijmegen Medical Center, University Medical Center Groningen, Accare, and VU University Amsterdam. The research leading to these results also received funding from the European Community’s Seventh Framework Programme (FP7/2007– 2013) under grant agreement number 278948 (TACTICS) and number 602450 (IMAGEMEND). Dr. Franke is supported by a Vici grant from NWO (grant number 016-130-669). The authors acknowledge the Department of Pediatrics of the VU University Medical Center for having the opportunity to use the mock scanner for preparation of our participants.

**NIDA**: Supported by the Intramural Research Programs of the National Institute on Drug Abuse and National Institute on Alcohol Abuse and Alcoholism

**TAOS**: The TAOS study (PI DEW) was supported by the National Institute on Alcohol Abuse and Alcoholism (R01AA016274) - “Affective and Neurobiological Predictors of Adolescent Onset AUD” and the Dielmann Family. The authors would like to acknowledge the TAOS staff and families and the Department of Psychiatry at the University of Texas Health Science Center at San Antonio that made this work possible.

**QTIM:** The QTIM study was supported by the National Health and Medical Research Council, Australia (Project Grants No. 496682 and 1009064 to MJ Wright) and the National Institutes of Child Health and Human Development (R01HD050735 to MJ Wright). MRI acquisition was undertaken at the Wesley Hospital and Centre for Advanced Imaging, University of Queensland, Brisbane and analyses at the Imaging Genetics Center, Keck School of Medicine of the University of Southern California. We thank the twins for their generosity of time and willingness to participate in our studies, the many radiographers and research assistants who collected the data, and the several research-associates who contributed to various aspects of the analyses.

